# The Gradient Clusteron: A model neuron that learns via dendritic nonlinearities, structural plasticity, and gradient descent

**DOI:** 10.1101/2020.12.15.417790

**Authors:** Toviah Moldwin, Menachem Kalmenson, Idan Segev

## Abstract

Synaptic clustering on neuronal dendrites has been hypothesized to play an important role in implementing pattern recognition. Neighboring synapses on a dendritic branch can interact in a synergistic, cooperative manner via the nonlinear voltage-dependence of NMDA receptors. Inspired by the NMDA receptor, the single-branch clusteron learning algorithm (Mel 1991) takes advantage of location-dependent multiplicative nonlinearities to solve classification tasks by randomly shuffling the locations of “under-performing” synapses on a model dendrite during learning (“structural plasticity”), eventually resulting in synapses with correlated activity being placed next to each other on the dendrite. We propose an alternative model, the gradient clusteron, or G-clusteron, which uses an analytically-derived gradient descent rule where synapses are “attracted to” or “repelled from” each other in an input- and location- dependent manner. We demonstrate the classification ability of this algorithm by testing it on the MNIST handwritten digit dataset and show that, when using a softmax activation function, the accuracy of the G-clusteron on the All-vs-All MNIST task (∼85%) approaches that of logistic regression (∼93%). In addition to the location update rule, we also derive a learning rule for the synaptic weights of the G-clusteron (“functional plasticity”) and show that a G-clusteron that utilizes the weight update rule can achieve ∼89% accuracy on the MNIST task. We also show that a G-clusteron with both the weight and location update rules can learn to solve the XOR problem from arbitrary initial conditions.

## Introduction

In the discipline of machine learning, artificial neural networks (ANNs) have gained great popularity due to their impressive success in solving a wide variety of computational tasks. ANNs also hold a special appeal due to their similarity to the networks of neurons in biological brains, lending hope to the possibility that ANNs might serve as a general-purpose algorithm for replicating human-like intelligence. ANNs have also been used by computational neuroscientists to simulate activity dynamics and learning processes in the brain.

Despite their utility for both machine learning and neuroscience, ANNs operate at a level of abstraction that disregards many of the details of biological neural networks. In particular, the artificial neurons in ANNs integrate their inputs linearly; that is to say, each input to a neuron in an ANN is given an independent weight and the neuron applies an activation function to the linear weighted sum of these inputs.

In contrast, synapses in biological neurons display an assortment of nonlinear interactions due to the passive biophysical properties of the cell as well as the active properties of voltage-gated ion channels. One such nonlinear channel of particular interest is linked to the N-methyl-D-aspartate (NMDA) receptor. The NMDA channel is both ligand-gated and voltage-gated (Jahr and Stevens 1990; Mayer, Westbrook, and Guthrie 1984; Nowak, Bregestovski, and Ascher 1984), allowing neighboring synapses on a dendrite to interact in a cooperative, supralinear manner. When two nearby NMDA-based synapses are activated simultaneously, the voltage-dependence of the NMDA receptor creates a voltage perturbation (depolarization) that is greater than the linear sum of the two inputs had they been activated independently (Polsky, Mel, and Schiller 2004).

The NMDA receptor has been shown to play a crucial role in structural plasticity, or the growth and elimination of synapses between neurons (Breton-Provencher, Coté, and Saghatelyan 2014; Kleindienst et al. 2011; Niculescu et al. 2018; Takahashi et al. 2012). Structural plasticity plays a crucial role in development and learning (Caroni, Donato, and Muller 2012; Elston and Fujita 2014; Trachtenberg et al. 2002; Yang, Pan, and Gan 2009). Structural plasticity increases the likelihood that temporally correlated synapses are placed next to each other (El-Boustani et al. 2018; Kleindienst et al. 2011; Lu and Zuo 2017; Takahashi et al. 2012; Winnubst et al. 2015), which may take advantage of the NMDA supralinearity. While some structural plasticity may be due to new presynaptic-postsynaptic pairs being formed, structural plasticity can allow neurons with extant synaptic connections between them to find the most functionally effective location to make a synapse. The commonality of multiple synaptic contacts between a single presynaptic-postsynaptic pair of cells, as shown by electron microscopy (Bartol et al. 2015; Fares and Stepanyants 2009; Kasthuri et al. 2015; Motta et al. 2019), as well as evidence that the number of multiple synapse boutons increases in an enriched environment (Jones et al. 1997) lends support to the idea that connected neurons may be sampling dendritic locations so as to form optimal functional clusters.

The synergistic coactivation of neighboring inputs on the dendrite via nonlinear NMDA receptors and the ability of a neuron to relocalize its synapses in response to correlated activity have been theorized to have a computational function. An early framework (Mel 1991, 1992) proposed that, by placing synapses with correlated inputs next to each other, a neuron could learn to solve a classification task. To demonstrate this capability, Mel created a simplified model of a neuron consisting of a single long dendrite, called the *clusteron*. The dendrite consisted of a sequence of discrete locations, from 1 to N, where N is the number of features in the input (in an image, N would be the number of pixels). Each input to the neuron would impinge upon a specific dendritic location. The “activation” of each synapse was defined as the direct input to that synapse multiplied by the sum of the inputs to nearby synapses within a fixed radius on the dendrite.

In order to train the clusteron to recognize a class of patterns (e.g. to identify pictures of dogs), the clusteron is presented with patterns from the class to be recognized (called the positive class) and stores the average activation of each synapse per pattern. At every epoch of learning, the clusteron randomly swaps the locations of “poorly performing” synapses (i.e. synapses with a low activation relative to the other synapses) with each other on the dendrite, eventually resulting in a configuration wherein correlated inputs become spatially adjacent to each other. The adjacent correlated synapses interact nonlinearly with each other, leading to a higher overall activation for the positive class of patterns.

In this work, we propose an alternative model for learning via dendritic nonlinearities and structural plasticity on the single dendrite, which we call the *gradient clusteron*, or G-clusteron. Unlike the original clusteron model, the G-clusteron uses a dendrite with continuous-valued locations (as opposed to the discrete locations in the clusteron) and a bell-shaped distance-dependent function to model the location dependence of interactions between synapses. These modifications allow us to analytically derive a gradient descent-based learning rule for the synaptic locations of the inputs to the G-clusteron. We show that the G-clusteron’s location-update rule can learn to solve the classic MNIST handwritten digit multiclass classification task with accuracy comparable to a linear classifier (logistic regression). Moreover, we derive an additional plasticity rule for the synaptic weights of the G-clusteron and show that when the synaptic location rule and the synaptic weight rule are used simultaneously, the G-clusteron can learn to solve the exclusive or (XOR) binary classification task, a feat which cannot be accomplished by a linear classifier (Minsky and Papert 1969).

## Results

The G-clusteron is a model neuron with a single one-dimensional “dendrite” containing synapses at various dendritic locations (Figure 1A). The *activation* of a synapse is defined as the product of its weighted input with a distance-weighted sum of the weighted input of every synapse on the dendrite (including itself). The G-clusteron then applies a nonlinearity to the sum of its synaptic activations.

**Figure 1.**
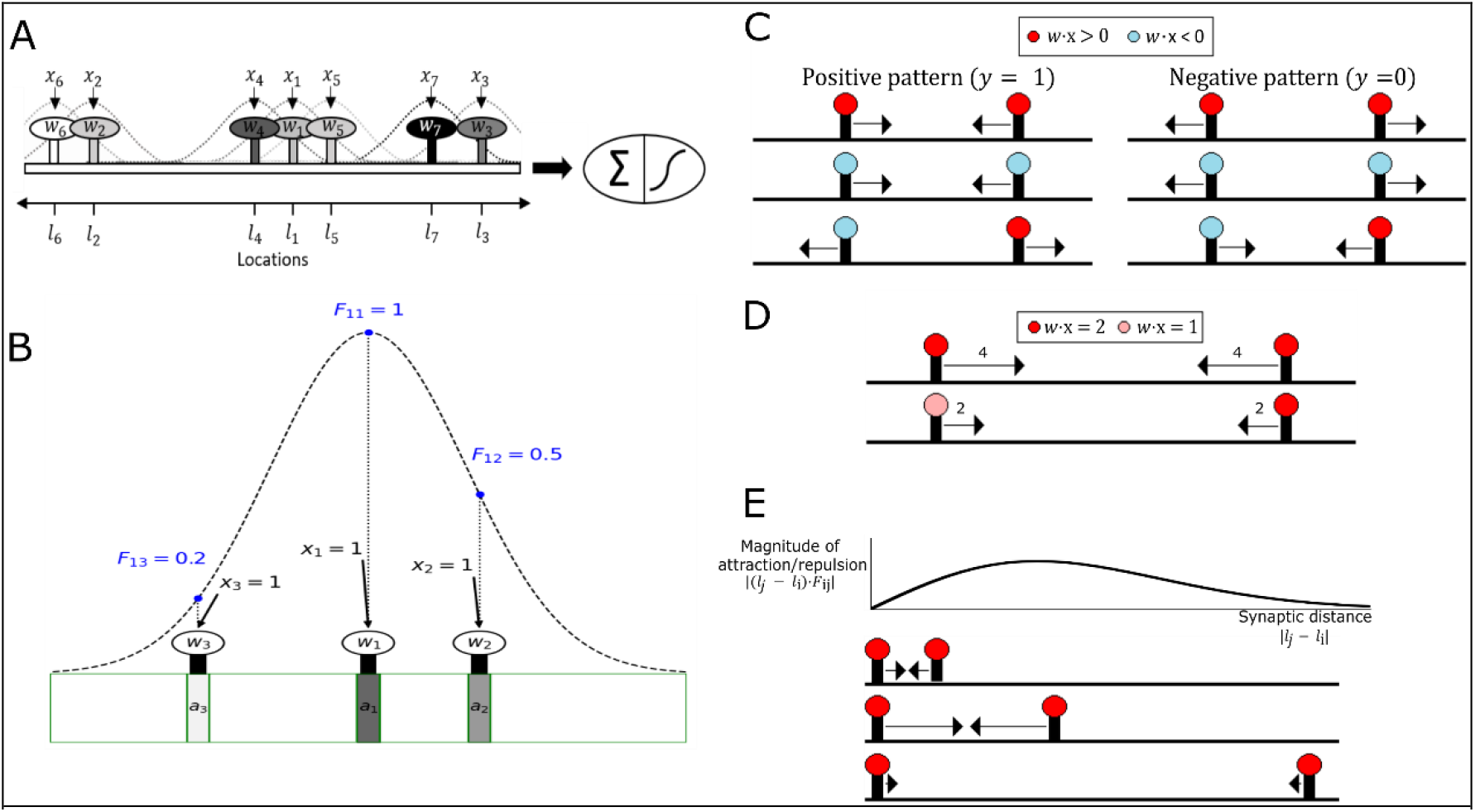
G-clusteron model and synaptic location update rule. (A) Schematic of the G-clusteron. Each input *x*_*i*_ is associated with a synapse at location *l*_*i*_ and weight *w*_*i*_. The activation of each synapse is affected by each other synapse according to a distance-dependent factor (dashed curves). A threshold function is applied to the sum of the activations. (B) Schematic of distance-dependent interactions [Eqs. (2) and (3)[. The distance-dependent factor between synapse 1 and synapses 2, 3, and itself are shown as blue points along the distance-dependent function (dashed curve). Synapse 1 interacts maximally with itself, moderately with synapse 2, and slightly with synapse 3. Even though the inputs and weights for all synapses are identical, the activations (shaded rectangles, darker shades indicate larger activation) differ due to the relative locations of the synapses (e.g. synapse 1 is centrally located so it is affected by synapses 2 and 3 more than synapses 2 and 3 affect each other). (C) Schematic of the sign-dependence of the location update rule (Eq. 7). Colors of the synapses indicate the sign of the weighted inputs. For positive patterns (left column), two synapses attract each other if their weighted inputs have the same sign (left column, top and center) and repel each other if the weighted inputs have opposite signs (left column, bottom). In negative patterns this is reversed (right column). (D) Effect of input and weight magnitude on the magnitude of the location update rule. The magnitude of the attraction and repulsion (arrow length) is proportional to |*w*_*i*_*x*_*i*_ ∗ *w*_*j*_*x*_*j*_|. (E) Top: Magnitude of the location update rule as a function of the distance between two synapses. Bottom: Three examples for the location-dependent update rule for positive input pattern (aligned to the curve above). Arrow length denotes magnitude of update. This magnitude is multiplied by the weighted input factor of the learning rule from (D) as well as the magnitude of the error | *ŷ* – *y* | to obtain the final magnitude.

Formally, for a given input pattern ***x*** = [*x*_1_, *x*_2_ … x_*N*_], the activation *a*_*i*_ of synapse *i* is:

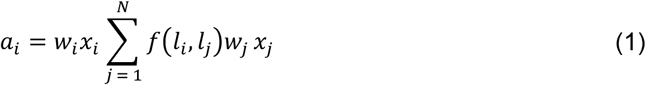

Where *l*_*i*_ is the real-valued location of synapse i on the dendrite, *w*_*i*_ is the weight of synapse i, and f(*l*_*i*_, *l*_*j*_) is a bell curve-shaped distance-dependent function which determines how much each synapse affects each other synapse, defined as:

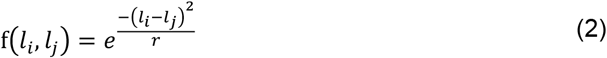

where *r* is a hyperparameter that determines the width of the curve. (From a biophysical standpoint, Eq. 2 is analogous to the distance-dependence of the voltage response of a cable to an instantaneous current impulse (Koch 2004). In this framework, *r* can be thought of as the square of the cable length constant *λ*, where 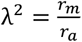, or the ratio of the membrane resistance to the axial resistance.)

Note that f(*l*_*i*_, *l*_*j*_) = 1 when *l*_*i*_ = *l*_*j*_ (i.e. when synapses i and j occupy the same location) and that f(*l*_*i*_, *l*_*j*_) approaches 0 as synapses *i* and *j* move further away from each other (Figure 1B). Also note that f(*l*_*i*_, *l*_*j*_) = f(*l*_*j*_, *l*_*i*_). For convenience of notation and computation, we define a matrix F such that

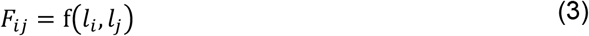

F is thus a symmetric matrix with ones on its diagonal.

For a given pattern of presynaptic inputs ***x***, the G-clusteron sums its activations together with a bias term b to produce:

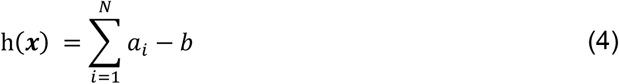

Because we will be using the G-clusteron as a binary classifier, we apply a sigmoidal nonlinearity g and produce an output *ŷ* between 0 and 1:

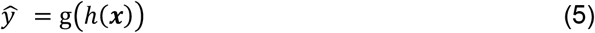

As in logistic regression, *ŷ* can be interpreted as a probability estimate for the binary label for the input pattern ***x***, with values closer to 0 representing a prediction for the label 0 (called the negative class) and values closer to 1 representing a prediction for the label 1 (called the positive class).

The G-clusteron thus differs from the original clusteron (Mel 1991) in two important ways: 1) each synapse has a real-valued location on the dendrite instead of an integer-valued location index and 2) each synapse’s activation function depends on the inputs of its neighbors as a gradually decreasing distance-dependent function as opposed to a hard cutoff at a fixed distance.

Defining the output of the G-clusteron in this fashion allows us to derive a gradient descent plasticity rule for the synaptic locations (see Methods). For each input pattern presented to the G-clusteron, we update each synaptic location according to the rule:

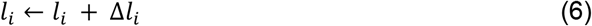

where:

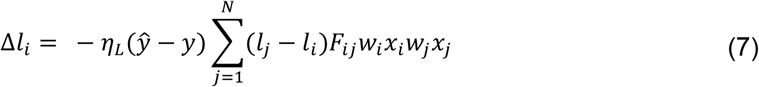

where y ∈ 0,1 is the true label (negative or positive class) for pattern ***x***, and *η*_*L*_ is the learning rate for the synaptic locations. This gradient rule for each synapse can be understood as summing over “forces” that depend on pairwise interactions between that synapse and each of the other synapses on the dendrite. The interactions between two synapses depend both on the weighted inputs of the synapses and the distance between the synapses in the following manner:

1. For positive-class training patterns, synapses with same-sign weighted inputs exhibit “attraction” while synapses with opposite-sign weighted inputs exhibit “repulsion” (Figure 1C, left).
2. For negative-class training patterns this trend is reversed: same-sign synapses are repelled, while opposite-sign synapses attract (Figure 1C, right).
3. The magnitude of attraction and repulsion between two synapses is proportional to the product of the weighted inputs of the two synapses (Figure 1D).
4. The magnitude of attraction and repulsion between two synapses is distance-dependent. The attractive/repulsive force is small at very small distances, becomes larger at intermediate distances, and shrinks again at large distances (Figure 1E).

(The magnitude of the update for each synapse is also scaled by the magnitude of the error term, |*ŷ* − *y* |).

Eq. 7 can be interpreted as a vector field along the dendrite, like force fields in physical systems of particles. If we consider a “unit synapse” such that *wx* = 1 at an arbitrary location *l* on the dendrite, the magnitude and direction of the “force vector” created by the plasticity rule at that location for a given input pattern ***x*** is given by:

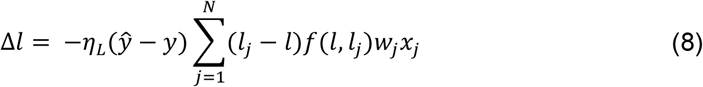

This interpretation of the location plasticity rule can give us some intuition as to how such an algorithm might be implemented biologically. We can imagine extracellular chemical signals being released at regions of the dendrite where there is a concentration of synaptic activity. Excitatory and inhibitory inputs would have different chemical signals associated with them. The diffusion of these chemicals in the extracellular space around the dendrite would create a location-dependent field of chemical gradients. These chemical gradients could induce presynaptic axons to form or eliminate synapses on the dendritic regions that were recently active, with differential effects for excitatory and inhibitory synapses.

### Toy Examples

To illustrate how the location update learning rule works, we consider several toy problems. For these problems, instead of training the G-clusteron to discriminate between two classes by presenting patterns from the positive and negative classes in a random order, we will show what happens when the G-clusteron is given a dataset where all examples are from the positive class (in other words, the G-clusteron is tasked with maximizing its output on all the input patterns, see Methods) and when all examples are from the negative class (so the G-clusteron must minimize its output for all input patterns). [For all of the following examples, all synaptic weights are fixed to have the value 1 and don’t change over the course of learning; our results thus strictly depend on the synaptic locations and inputs].

For the first toy problem, we create a single synaptic input vector ***x*** where the values of ***x*** are randomly distributed between -1 and 1. This input vector is repeatedly presented to the G-clusteron with a positive label (y = 1), such that the G-clusteron will attempt to maximize its overall activation. Because the interaction between synapses in Eq. 1 is multiplicative, a sensible strategy would be to minimize the distance between synapses with same-sign inputs and maximize the distance between synapses with opposite-sign inputs. This would result in two clusters on the dendrite: one with positive-valued synaptic inputs and one with negative-valued synaptic inputs. The location update rule, by causing attraction between same-sign synapses and repulsion between opposite-sign synapses, does exactly this (Figure 2A, S1 Movie).

**Figure 2.**
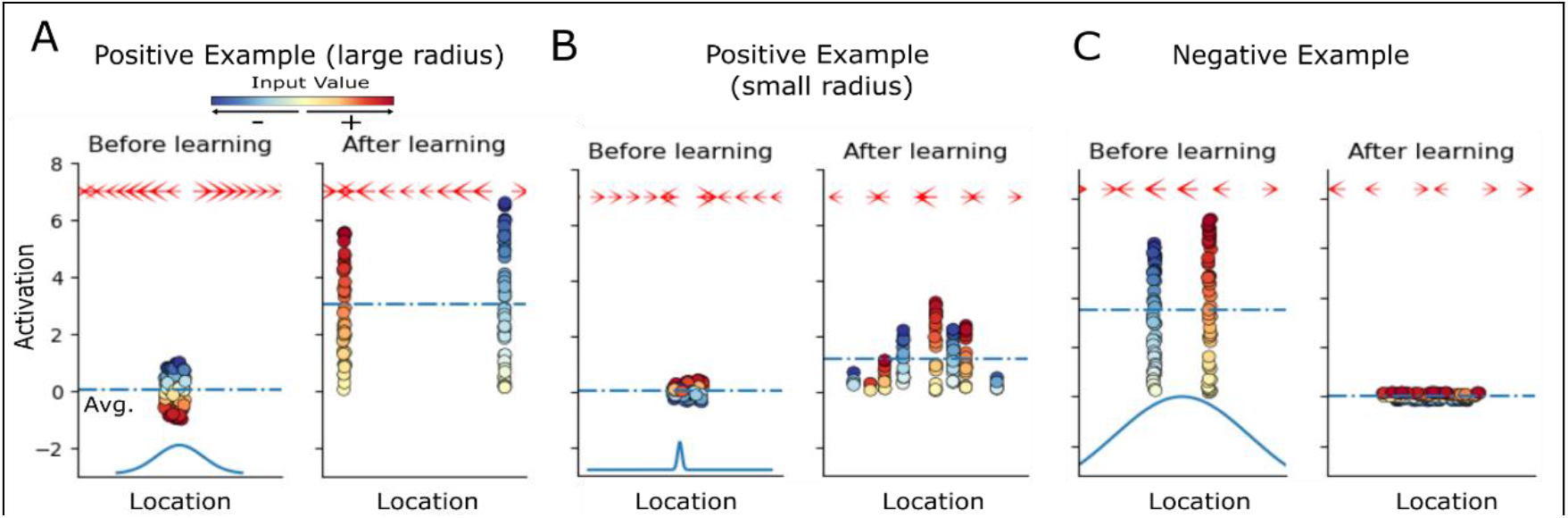
Dynamics of synaptic clustering due to the G-clusteron update rule. (A) Training the G-clusteron on a single positive pattern (see S1 Movie). Synaptic input values (colored circles) were drawn uniformly between -1 and +1 (all weights are fixed at 1 for the duration of the example). Position of circles along the X-axis denotes synaptic location on dendrite; their location along the Y-axis denotes synaptic activation value. In the initial epoch (left), synaptic locations are randomly initialized in close proximity. During learning, synapses with positive inputs move in the opposite direction from synapses with negative inputs. In the final epoch (right) separate clusters are observed for positive and negative inputs. Dashed blue line shows the average synaptic activation, which in this case is increased by the plasticity rule. Arrows at the top of each panel denote the magnitude and direction of the “force field” created by the plasticity rule (Eq. 8); by convention, the arrow at each location points in the direction that a unitary positive input (*wx* = 1) would move according to the plasticity rule. Blue curve at the bottom of the left panel shows the width of the distance-dependent function (dependent on the hyperparameter *r*) for this example. (B) Same as A with a smaller value for *r* (see S2 Movie). Here the plasticity rule operates on a more local scale, creating multiple smaller same-sign clusters instead of two large clusters as in A. (C) Same as A for a negative pattern (see S3 Movie). Before learning, synapses are initialized in separate clusters for positive and negative inputs (left). Over the course of the learning, synapses are pulled toward each other until they are intermixed (right). Note that after learning the average activation level decreases.

We now show how the hyperparameter *r* (i.e. the width of the distance-dependent curve) affects the learning rule for input patterns from the positive class. In the first example, r was sufficiently large relative to the initial dispersion of synapses such that all of the synapses could “feel” each other, thus enabling all synapses with the same sign to eventually aggregate into a single cluster. However, if r is reduced, synapses that start further apart from each other do not exert a strong pull on each other, so instead the learning rule operates in a more local fashion, creating several same-sign clusters (Figure 2B, S2 Movie).

We next train the G-clusteron on the same input pattern, but this time with a negative label (y = 0), such that the G-clusteron will minimize its overall activation. Here we pre-initialize the G-clusteron in a high-activation state where same-sign synapses start in separate clusters. Inversely to the first strategy, we would like opposite-sign inputs to become intermixed. Because the plasticity rule flips attraction and repulsion for input patterns rom the negative class, we achieve the expected result (Figure 2C, S3 Movie).

Having demonstrated that the G-clusteron learns by aggregating and dispersing same-sign and opposite-sign synaptic inputs for individual input patterns, we illustrate that this mechanism serves to learn the correlation structure of a dataset comprised of multiple input patterns. To this end, we create a 20-dimensional multivariate Gaussian distribution where each dimension of the Gaussian had a mean of 0 and a covariance matrix Σ such that:

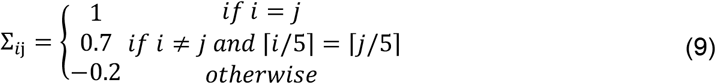

In other words, dimensions *x*_1_ … *x*_5_ were all correlated with each other, as were dimensions *x*_6_ … *x*_10_, *x*_11_ … *x*_15_, and *x*_15_ … *x*_20_, but otherwise, dimensions were negatively correlated (Figure 3A). As such, each vector sampled from this distribution would exhibit similarity between the first five dimensions, second five dimensions, and so forth (Figure 3B).

**Figure 3.**
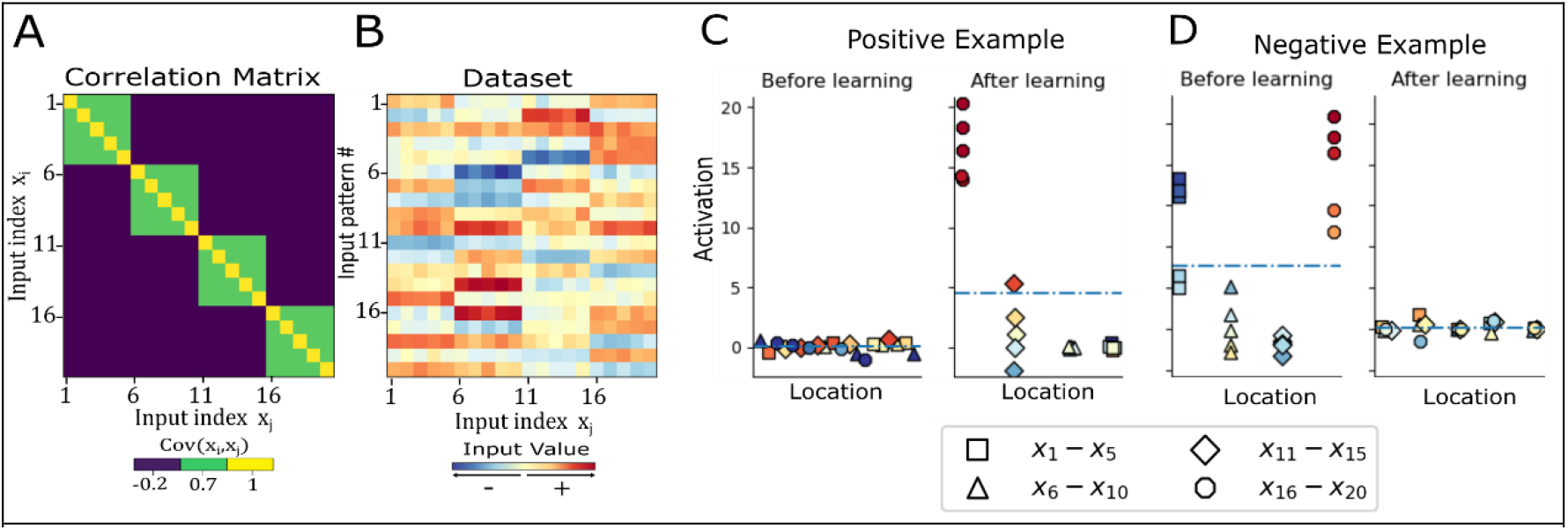
Dynamics of synaptic clustering for a multivariate Gaussian dataset with correlated input dimensions. (A) Covariance matrix for multivariate Gaussian example (Eq. 9). This covariance matrix generates samples where dimensions *x*_1_ … *x*_5_, *x*_6_ … *x*_10_, and so on are positively correlated with each other, while all other correlations are negative. (B) Patterns drawn from the multivariate Gaussian defined with the covariance matrix in (A). Each row is an input pattern presented to the G-clusteron. (C) The G-clusteron is initialized (left) with randomized synaptic locations and presented with positive-labeled input patterns from (B) (see S4 Movie). Correlated inputs have the same shape; colors indicate input values for the pattern presented in the epoch as in B. To increase activation, the G-clusteron groups together correlated inputs (right). (D) Same as (C) but for a negative example (see S Movie). For illustration, the G-clusteron is initialized with correlated inputs clustered together, as in the right panel of (C). Over the course of learning the synapses eventually form clusters of negatively correlated synapses.

As the mean of each dimension is 0, a linear neuron would be unable to increase its average output on this dataset, as there is no independent information in each synapse that could be given a linear weight. However, the G-clusteron can take advantage of the correlational structure of the data. Because the groups of positively correlated inputs will tend to all be either all-positive or all-negative in any given input pattern, the learning rule will gradually move the positively correlated synapses together, creating dendritic clusters of synapses with correlated inputs. As these clusters form, the output of the G-clusteron increases (Figure 3C, S4 Movie). Inversely, the G-clusteron can learn to decrease its output on this dataset by clustering negatively correlated inputs (Figure 3D, S5 Movie).

### Learning MNIST

The ability of the G-clusteron to learn the correlational structure of a dataset enables it to perform supervised classification by aggregating correlated inputs from the patterns where y = 1 and disaggregating correlated inputs from the patterns where y = 0 (Figure 4A). We trained the G-clusteron on the MNIST dataset of images of handwritten digits (Figure 4B) and compared its performance to that of the original clusteron from (Mel 1991) as well as to logistic regression. Before attempting the all-vs-all classification task, we test the G-clusteron separately on each digit in a one-vs-all classification paradigm. For each digit from 0-9, a G-clusteron was trained on a dataset where half of the images were of that digit (positive class, label y = 1) and half of the images contained other digits (negative class, label y = 0). The G-clusteron was then tested on a holdout test set (See Methods). This procedure was repeated for the original clusteron (only positive-class training examples are used for the original clusteron, see Methods) as well as logistic regression. The learning and testing process was repeated 10 times for each classifier to ensure performance stability. The results are shown in Table 1.

**Table 1.** Accuracies of one-vs-all on the MNIST dataset. Values are averaged over ten runs, with standard deviations in parentheses.

|  | 0 | 1 | 2 | 3 | 4 | 5 | 6 | 7 | 8 | 9 |
| --- | --- | --- | --- | --- | --- | --- | --- | --- | --- | --- |
| Clusteron | 92.8<br>(0.8) | 83.6<br>(5.3) | 82.5<br>(0.5) | 85.0<br>(0.5) | 77.6<br>(1.7) | 74.4<br>(1.5) | 87.5<br>(0.8) | 87.8<br>(0.8) | 80.2<br>(1.1) | 81.3<br>(1.0) |
| Gradient Clusteron | 96.8<br>(0.4) | 93.9<br>(4.4) | 87.6<br>(5.3) | 88.5<br>(0.9) | 91.8<br>(1.1) | 80.7<br>(4.1) | 95.1<br>(0.4) | 90.1<br>(5.0) | 82.3<br>(4.3) | 85.8<br>(3.6) |
| Logistic Regression | 98.2<br>(0.0) | 98.3<br>(0.0) | 94.4<br>(0.0) | 93.3<br>(0.0) | 96.7<br>(0.0) | 91.6<br>(0.0) | 96.5<br>(0.0) | 95.4<br>(0.0) | 91.2<br>(0.0) | 92.3<br>(0.0) |

**Figure 4.**
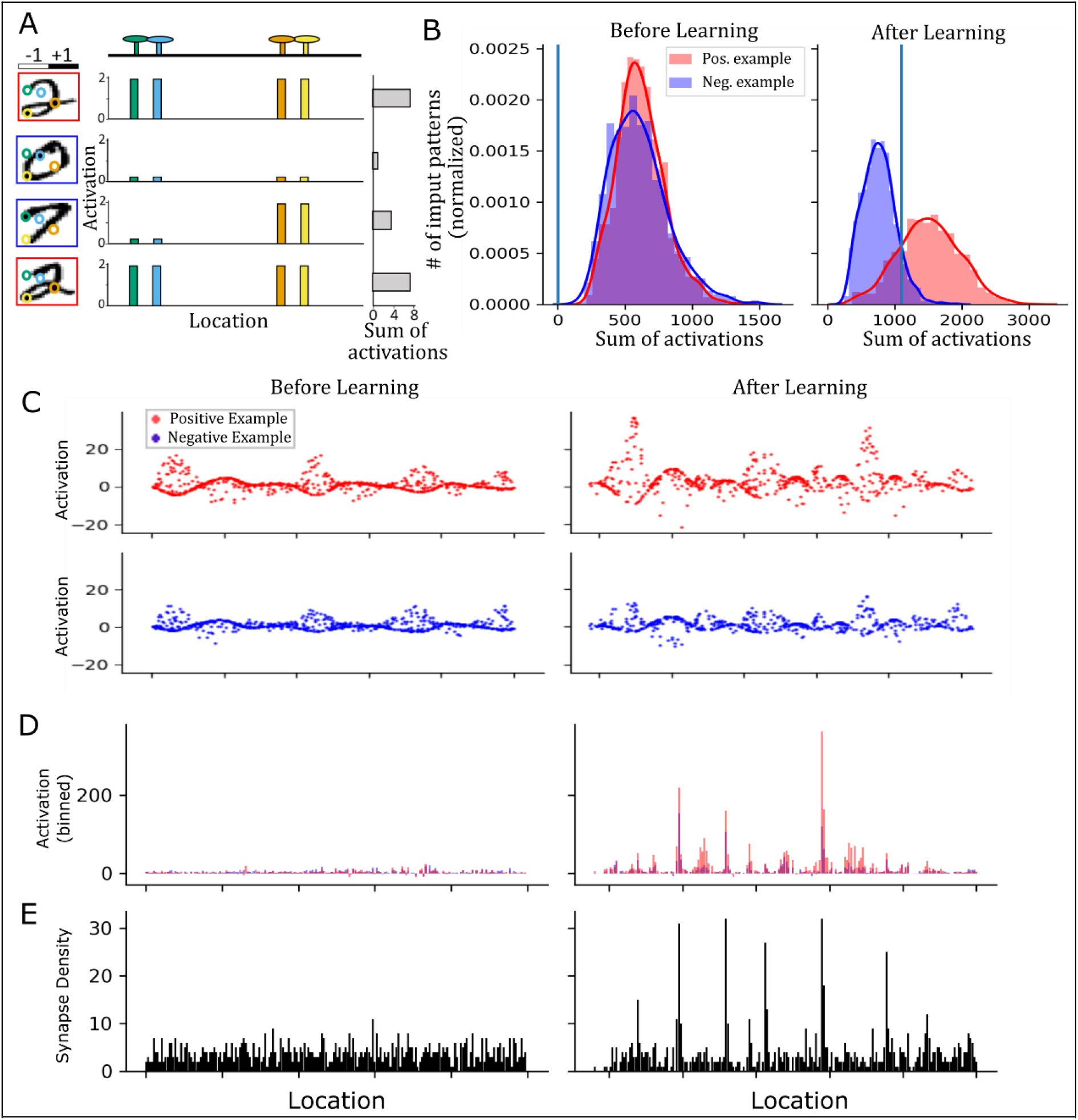
G-clusteron classification of handwritten digits (MNIST) and the resultant distribution of synaptic locations and activations. (A) Schematic of G-clusteron performing the one-vs-all classification task, where it must produce a larger activation for images containing the positive class 2 (digit with red border) than any other number (digit with blue border). Colored circles in the digit images indicate exemplar pixel locations mapped to exemplar synapses on the dendrite with the same color, generating two dendritic clusters (green-blue and orange-yellow, respectively). The synapses within these clusters will be maximally activated if both inputs to the cluster are the same sign (i.e. both black or both white), while opposite sign inputs will result in lower activation. (B) Exemplar histogram of sum of synaptic activations from the G-clusteron tasked with classifying the digit 2 (label *y* = 1, red) vs. all other digits (label *y* = 0, blue) before learning (left) and after learning (right). Over the course of learning, the G-clusteron increases its activation on images containing a 2 relative to images with other digits, enabling binary classification. Blue vertical line is the value of the bias term. (C) Synaptic activations for negative (top) and positive (bottom) patterns before (left) and after (right) learning. Note the increase in synaptic activations for positive patterns during learning. (D) Sum of the activations within small bins of dendritic length averaged over positive (red) and negative (blue) datasets before and after learning (aligned with (C)). Note that there are small regions (functional clusters) where the activations for positive patterns (red) is much larger than for negative patterns (blue). (E) Synaptic density per bin (same for positive and negative input patterns, aligned with (C-D)). Note that there are several high density regions (structural clusters) that sometimes correspond to the functional clusters.

For all digits, all three classifiers achieved a classification accuracy far above chance level of 50%. Depending on the particular digit, the clusteron achieved between 74.4% and 92.8% accuracy, the G-clusteron achieved between 80.7% and 96.8% accuracy, and logistic regression achieved between 91.2% and 98.3% accuracy. For each digit, the G-clusteron outperformed the clusteron, and logistic regression outperformed the G-clusteron. We emphasize that both the clusteron and the G-clusteron had all their weights fixed to 1 during the entire learning procedure and were only able to update their synaptic locations. Our results on the one-vs-all MNIST task thus demonstrate the remarkable efficacy of structural plasticity-based learning and suggests that the G-clusteron algorithm may be superior to the original clusteron algorithm for solving certain tasks.

The distribution of synapses and synaptic activations before and after learning MNIST (Figure 4C-E) can be instructive in understanding how the G-clusteron operates. In many instances, over the course of learning, the synapses on the dendrite moved from an approximately uniform synaptic density to a more clustered structure, with some regions of the dendrite having a higher synaptic density than others (Figure 4E). These higher density areas occasionally had larger activations for patterns from the positive class than for patterns from the negative class (Figure 4D), suggesting that the G-clusteron may be building structural clusters to take advantage of correlated inputs in the positive class relative to the negative class. However, not all high-density clusters showed high activation, and some high-activation regions were low density. Thus, while the G-clusteron learning algorithm does produce structural clusters as a consequence of learning, there is not a guaranteed correspondence between the structural clusters and activation level in a complex task like image classification.

Having shown that the clusteron and G-clusteron exhibit satisfactory performance on the one-vs-all MNIST task, we now turn to the harder problem of all-vs-all classification. Here, we wish to train a single-layer network of 10 classifiers on the MNIST dataset (one for each digit) and have the network classify each digit correctly. We consider two standard ways to train a single-layer network on a multiclass classification problem. In the one-vs-rest (OVR) method, 10 units are independently trained on a one-vs-all paradigm, as before, and the classifier which produces the largest output (*ŷ*) for a given input pattern is declared the “winner”, and the input pattern is assigned to the positive class for that classifier (Figure 5A). In the softmax (SM) method, the units are trained simultaneously on each example and the raw outputs (*h*(*x*)) of all units are passed through a softmax function (see Methods), which normalizes the output of each unit by the sum of the outputs of all the units in the layer (Figure 5B). This normalized output is then used in the error term *ŷ* − *y* when calculating the update rule. Because the softmax method allows the classifiers to communicate with each other via the output normalization, it can often lead to superior results for multiclass classification (Duan et al. 2003).

**Figure 5.**
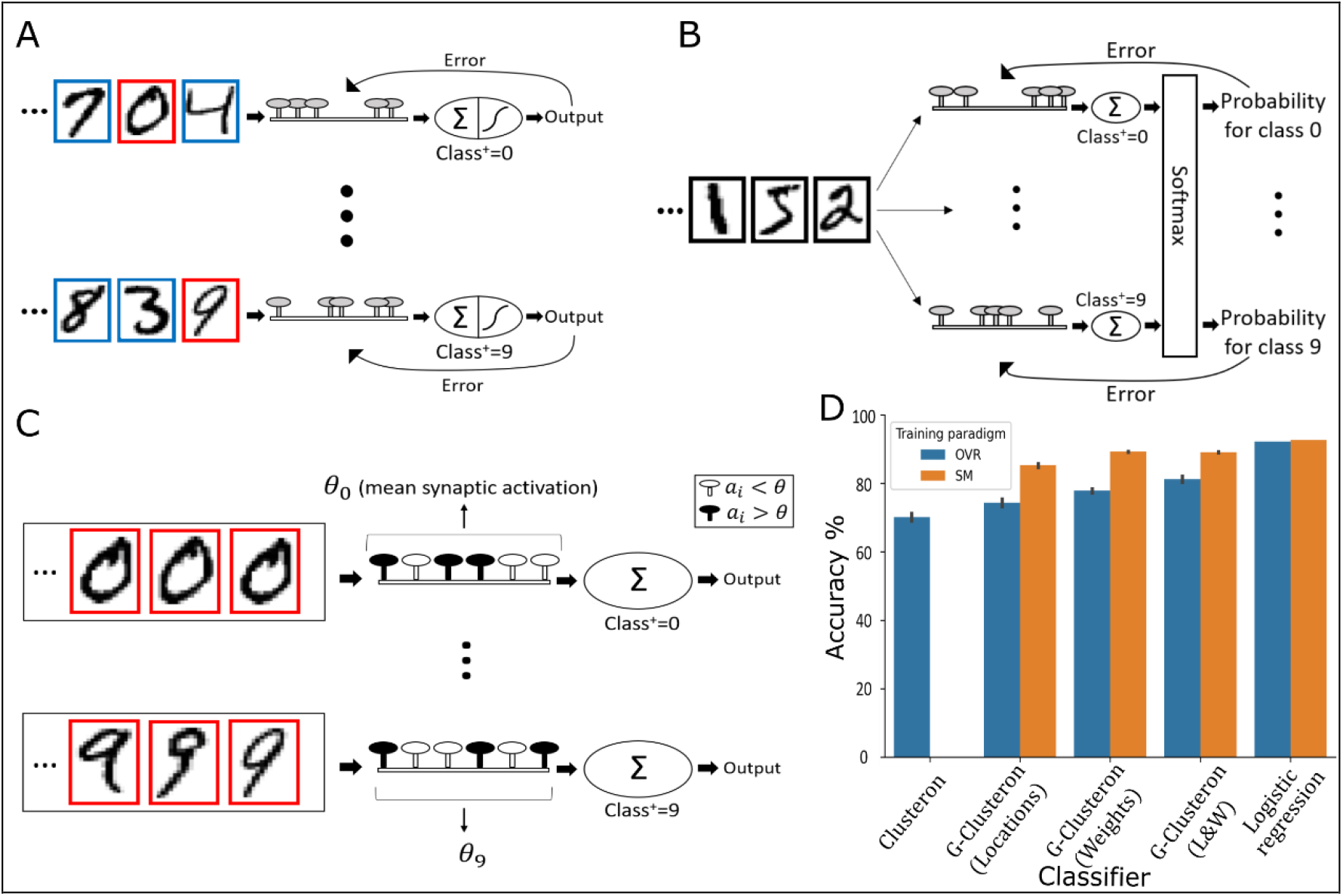
Multiclass classification with the G-clusteron on handwritten digits (MNIST) (A) One-vs-rest (OVR) learning scheme for the G-clusteron. Classifiers for each digit are trained independently on both positive (red border) and negative (blue border) digits. (B) Softmax (SM) classification scheme for the G-clusteron. Classifiers are trained simultaneously on each example and the sum of the synaptic activations for each classifier are fed into a softmax function which normalizes the output of each classifier by the output of all the classifiers. The error term used to update each classifier for an input digit thus has information about the output of all the classifiers for that digit. (C) OVR learning scheme for the original clusteron. On each epoch, each classifier is presented with an entire dataset consisting only of positive examples (red borders), and *θ* -- the mean synaptic activation over the dataset - - is calculated. The Clusteron learns by randomly shuffling the locations of synapses whose activation was less than *θ*. Because each classifier only trains on positive examples and because there is no error term that can be communicated to other classifiers, the original clusteron does not lend itself to a softmax architecture as in (B). (D) Mean accuracy for Clusteron, G-Clusteron (with either only the location update rule, only the weight update rule, or both rules), and Logistic Regression on the All-vs-All MNIST task using either the OVR method (blue) or SM method (orange). Error bars indicate standard deviation. Note that the original clusteron cannot use the SM method.

Importantly, the original single-dendrite clusteron does not utilize an error signal in its learning rule, so the original clusteron can only be trained on the multiclass task with the OVR paradigm (Figure 5C). Both logistic regression and the G-clusteron, however, do use an error term, making it straightforward to perform multiclass classification with both the OVR and SM methods.

To test the performance of the G-clusteron under both multiclass classification paradigms, we trained the original clusteron, the G-clusteron, and logistic regression using the OVR method, and the G-clusteron and logistic regression (but not the original clusteron) using the SM method. As with the previous task, learning for all classifiers was repeated 10 times to ensure stability.

When using the OVR paradigm, the original clusteron achieved an average accuracy of 70.1%, the G-clusteron achieves 74.3% accuracy, and logistic regression achieves 92.2% accuracy. While both the original clusteron and the G-clusteron achieve performance far better than chance level of 10%, neither of them do nearly as well as logistic regression. However, when we train the G-clusteron using the SM method, it achieves an impressive accuracy of 85.3%, closing in on the accuracy of logistic regression’s 92.6% accuracy in the SM scheme (Figure 5D). Although not superior to logistic regression, 85.3% accuracy on an all-vs-all MNIST classification task is notable for an algorithm that can only modify synaptic locations and not synaptic weights.

### Weight update rule for the G-clusteron

In addition to the location update rule, we also derive a gradient descent rule for the weights of the G-clusteron of the form:

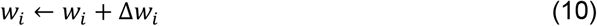

Where

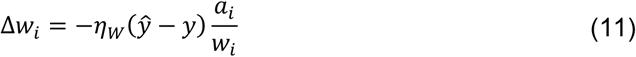

with ***η***_***W***_ as the learning rate for the weights. This rule can either be used on its own or in conjunction with the location update rule (Eq. 6) by simultaneously updating the weights and locations using their respective rules during each epoch. To see if the weight update rule can improve the accuracy of the G-clusteron on the all-vs-all MNIST task, we trained a single-layer G-clusteron network that used only the weight rule as well as a network that used both the location rule and the weight update rule simultaneously, using the same protocols (OVR and SM) as for the G-clusteron using only the location rule.

When only the weight rule was used, the G-clusteron achieved an accuracy of 77.9% for OVR and 89.3% for SM, and when the update rules were used simultaneously, the G-clusteron achieved improved an improved accuracy of 81.2% for OVR and a similar accuracy of 89.1% for SM. Thus, being able to manipulate synaptic weights may allow the G-clusteron to achieve slightly superior accuracy relative to a G-clusteron that can only update its synaptic locations (Figure 5D).

Although the G-clusteron did not perform as well as a linear classifier on the MNIST task, this does not indicate that the theoretical classification capacity of the G-clusteron is inferior to that of a linear classifier. In fact, we prove (see S1 Text) that if a G-clusteron with an arbitrary localization of synapses is equipped with a single additional “bias synapse” *x*_0_ (*x*_0_ = 1 for all input patterns, this is distinct from the 0^th^-order bias term *b*), it can approximate any linear classifier to an arbitrary degree of precision by appropriately assigning the weights. The discrepancy in classification accuracy in this task between the linear classifier and the G-clusteron with a weight update rule is thus likely due to the G-clusteron’s convergence to a local minimum rather than an inability to implement a linear separation boundary.

### XOR problem

To motivate the use of the weight update rule in combination with the location update rule for the G-clusteron, we consider the XOR function. The XOR function receives two bits of binary input and returns a 1 if the input bits are different or a 0 if the input bits are the same. The XOR function famously cannot be implemented by linear classifiers like the perceptron (Minsky and Papert 1969). It is thus valuable to demonstrate that the G-clusteron can implement this function. We first wish to show that there are values for the parameters (i.e. the weights and the locations of the two synapses) of the G-clusteron that will result in a correct implementation of the XOR function. We then show that a G-clusteron must be able to update both its synaptic locations and its synaptic weights if it is to learn to solve the XOR problem from an arbitrary initialization of its parameter values.

To implement the XOR function, the parameters of the G-clusteron must satisfy both of the following inequalities (See Methods):

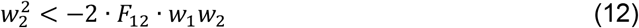

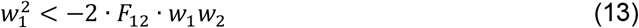

Note that although we originally have 4 parameters (*w*_1_, *w*_2_, *l*_1_, and *l*_2_), the solution space satisfying these inequalities (Figure 6A-B) can be expressed in terms of only three parameters: *w*_1_, *w*_2_, and *F*_12_.

**Figure 6.**
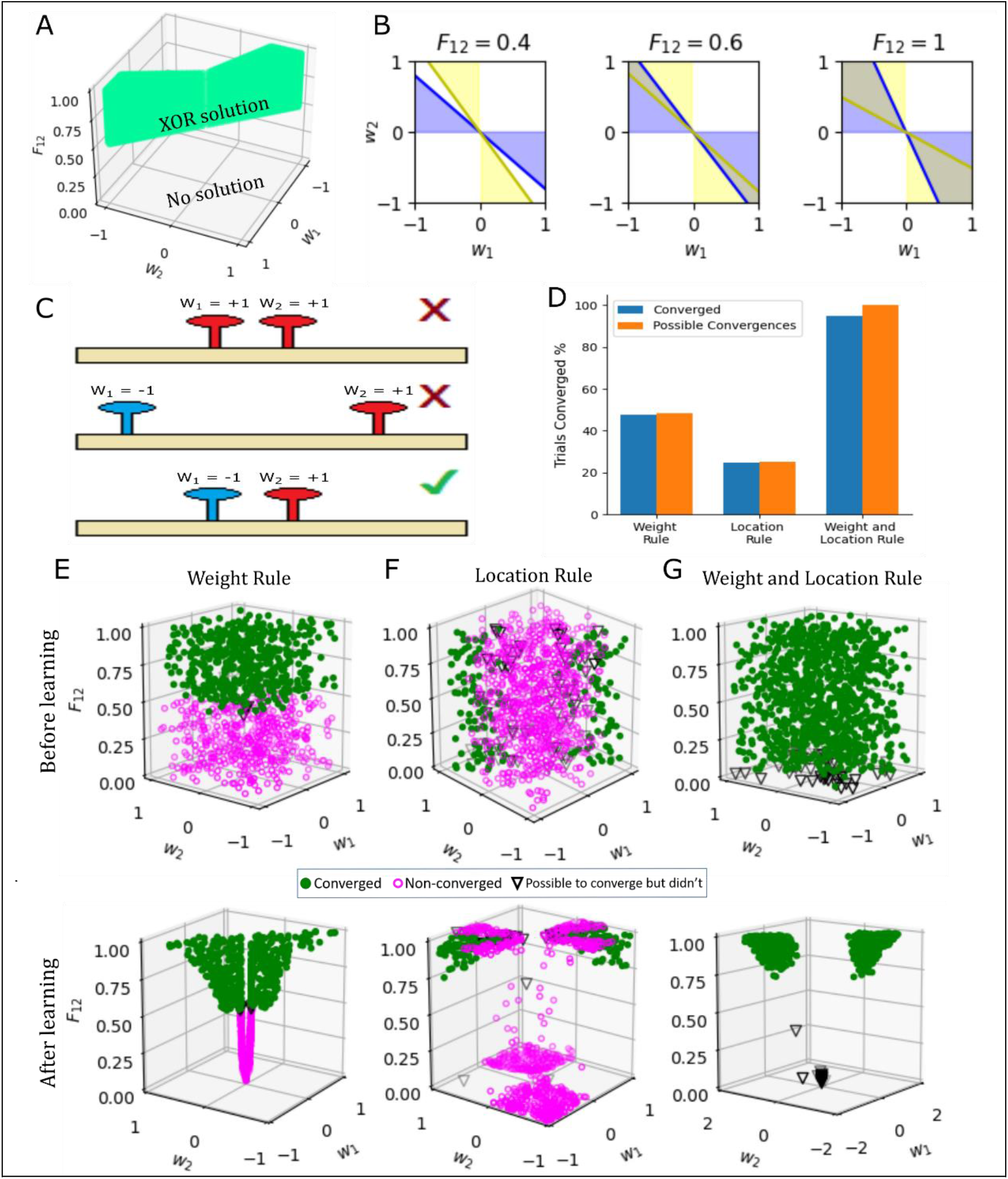
The G-clusteron can learn to solve the XOR problem from arbitrary initial conditions if and only if it can update both its locations and weights. (A) Analytical solution to XOR problem. Shaded area represents the parameter space in terms of the two weights, *w*_1_ and *w*_2_, and the distance-dependent factor, *F*_12_, which would produce the correct outputs for the XOR problem. (B) Representation in weight space of the analytical solution to XOR, shown for selected values of *F*_12_ (i.e. slices in the *F*_12_ axis of (A)). The blue region satisfies the inequality from eq. (12), and the yellow region satisfies the inequality from eq. (13). The G-clusteron can solve the XOR problem when both inequalities are satisfied (overlap region, same as colored region in A). Note that the solution space for the weights is null when *F*_12_ < 0.5 and increases as *F*_12_ approaches 1. However, there are many weight assignments that never satisfy both inequalities. (C) Examples of correct and incorrect parameter assignments for the XOR problem. Some weight assignments that never produce a correct solution for XOR regardless of the distance between the synapses (top). Moreover, if the synapses are too far away from each other, there are no weights that satisfy the XOR relation (middle). Thus, the solution for XOR requires that the synapses be close together and the weights appropriately set (bottom). (D) Number of trials that were analytically possible to converge vs. trials that successfully converged on the XOR problem for the G-clusteron trained with only the weight update rule, only the location update rule, or both rules. (E) Top: Convergence (marker shape and color, see legend) as a function of the initial parameters of 1000 randomly initialized G-clusterons with only the weight update rule. Note that only trials in which the initial value for *F*_12_ was greater than 0.5 converged. Bottom: Convergence as a function of the final parameters of the G-clusterons after learning. Note that converged solutions fall within the region specified in A. (F) Same as E for G-clusteron using only the location update rule. Note that only trials in which the initial values for the weights were within the overlap region of B (right panel) were able to converge. (G) Same as E for G-clusteron with both the weight and location update rules. Almost all solutions converge.

When expressed in this manner, there are several things to observe about these inequalities. First, note that for *F*_12_ ≤ 0.5, there is no way to assign the weights such that Eq. 12 and Eq. 13 are satisfied (Figure 6B, left). This means that if the two synapses are initialized sufficiently far away from each other such that their distance-dependent factor is less than 0.5, a G-clusteron with only a weight update rule (Eq. 10) would have no way to solve the XOR problem.

Moreover, the range of weights that are valid to solve the XOR problem increases as *F*_12_ increases from 0.5 to 1, where the solution space for the weights is maximal (Figure 6B, center and right). In other words, as the synapses move closer together, there is a larger set of weights that would satisfy Eq. 12 and Eq. 13. However, even when the synapses occupy the exact same location (i.e. *F*_12_ = 1), there is still a large range of invalid weights. For example, if both weights are initialized with the same sign, a G-clusteron with only a location update rule (Eq. 6) could not implement the XOR function (Figure 6B and 6C). Therefore, for the G-clusteron to solve XOR from arbitrary initial parameter values, it must be able to update both its weights and its locations.

To demonstrate this numerically, we created 3 G-clusterons: one with only a weight update rule, one with only a location update rule, and one with both update rules. For each G-clusteron, we ran 1000 trials with randomized initializations on the inputs and labels for the XOR problem. Each trial ran for 10,000 epochs or until convergence (see Methods).

A trial was considered possible to converge for a G-clusteron with only the weight update rule if there existed some weight assignment that would produce the correct XOR output given the initialization of the two synaptic locations, which would occur if the initial *F*_12_ was greater than 0.5. A trial was considered possible to converge for a G-clusteron with only the location update rule if there existed some location assignment that would produce the correct XOR output given the initialization of its synaptic weights. Because the solution space for XOR grows as the synapses move closer, this entails that the initial weights would have to satisfy Eq. 12 and Eq. 13 given that *F*_12_ = 1 (in other words, the weights must fall in the overlap shaded region of Figure 6B, right panel). For the G-clusteron with both the weight and the location update rules, all trials could potentially converge (Fig. 6D).

The G-clusteron with only the weight update rule converged for 475 trials out of 485 trials analytically possible to converge (Figure 6D, 6E), unexpectedly failing on trials where *F*_12_ was initialized slightly above the boundary condition of 0.5, because the solution space for the weights when *F*_12_ is fixed near 0.5 is small (Fig. 6B middle), and it can therefore take the learning algorithm an exorbitantly long time to direct the weights into this exactingly narrow range of values.

The G-clusteron with only the location update rule converged for 247 trials out of 251 trials analytically possible to converge (Figure 6D, 6F), unexpectedly failing on trials where *F*_12_ was initialized very close to 0, as before, and also on trials where the weights were initialized very close to the boundary of the solution space (Fig. 6B, right panel). In these cases, it was evidently difficult for the algorithm to find the bias value with sufficient precision to correctly perform the classification, even though the weights and locations were analytically valid.

Only the G-clusteron with both the weight and the location update rules was able to converge on almost all of the trials (947 out of 1000 trials), unexpectedly failing only on trials where *F*_12_ was initialized very close to 0 (Fig. 6D, 6G). These unexpected failures occurred because the magnitude of the location update rule for a given epoch scales with the value of *F*_12_ and the weights at that epoch (Eq. 7). Thus, if *F*_12_ is initialized very close to 0, the synaptic locations may only move a miniscule amount each time the gradient rule is applied, depending on the values of the other parameters.

## Discussion

We have shown that the G-clusteron is a robust single-neuron learning algorithm that can solve real-world classification tasks, such as MNIST handwritten digit classification. The G-clusteron’s ability to solve this task merely by moving synapses on its dendrite without using synaptic weights demonstrates the computational potential of structural plasticity that makes use of distance-dependent nonlinearities. For comparison, recent work to solve MNIST without training synaptic weights (via learning an effective network architecture) required a deep network with dozens of nodes per output class to achieve logistic regression-like performance (Gaier and Ha 2019). The G-clusteron achieves this level of accuracy using only a single neuron per output class (excluding the input layer).

While the original single-dendrite clusteron (Mel 1991) also exhibits impressive performance when tasked with one-vs-all classification on a single digit, the lack of a supervised gradient descent-based plasticity rule makes it difficult to scale this algorithm to a multiclass classification problem, as the clusteron learning rule does not have an error signal that can be communicated between neurons within a layer via a softmax activation function. [It should be noted that the multi-branch clusteron (Poirazi and Mel 2001) employs a supervised learning strategy that applies gradient descent to modify synaptic stability, which in turn determines how synapses are mapped to individual branches, however they do not directly learn synaptic locations via gradient descent as we do here.]

The gradient descent plasticity rule of the G-clusteron enables effective multiclass classification and allows the use of a variety of techniques developed for classic ANNs. For example, we used the ADAM momentum-based adaptive learning method (Kingma and Ba 2015) to dynamically optimize our learning rate. Bringing dendritic cluster-based plasticity algorithms within the fold of gradient descent learning opens up many possibilities for the extension of the G-clusteron algorithm using the extant mathematical frameworks and robust literature for learning using gradient descent in ANNs (Rumelhart, Hinton, and Williams 1986).

We have also shown that incorporating a plasticity rule for the weights in a G-clusteron alongside the dendritic location plasticity rule enables the G-clusteron to solve the classic linearly inseparable XOR problem (Minsky and Papert 1969). This reinforces the possibility that neuron models with dendritic nonlinearities may be more computationally powerful than the M&P nodes used in ANNs (Gidon et al. 2020; Poirazi, Brannon, and Mel 2003; Poirazi and Mel 1999, 2001).

### Relationship to Previous Work

Although our work here treats dendrites with NMDA receptors as nonlinear integration units, recent work (Moldwin and Segev 2020) has shown that a detailed biophysical model of a cortical layer 5 pyramidal cell with NMDA synapses and other active mechanisms can implement the perceptron learning algorithm, implying that a neuron can behave as a linear classifier. However, this does not preclude the possibility of nonlinear integration and plasticity rules; rather it should be taken as an indication a neuron can elect to apply a simple plasticity rule that ignores the synergistic interactions between synapses and still manage to solve classification tasks as a perceptron would.

The original clusteron (Mel 1991) fits within the framework of a subsampled quadratic classifiers, as the clusteron sums a subset of its *N*^2^ mixed terms *x*_*i*_*x*_*j*_ (Poirazi and Mel 1999). The G-clusteron can be thought of a “constrained” quadratic classifier in the sense that the equation for the G-clusteron contains all *N*^2^ mixed terms, but the coefficients of these terms – the entries of the *F* matrix – are “tied together” in the sense that *F* must be produced from a matrix representing distances between points in one dimension, and not all matrices are valid distance matrices. The G-clusteron therefore gives us *N*^2^ mixed terms for the price of *N* location parameters, but at the cost of not being able to manipulate each of the *N*^2^ coefficients independently.

In addition to the original clusteron and its multiple-layer extension (Poirazi, Brannon, and Mel 2003; Poirazi and Mel 2001), there have been a variety of other attempts to model learning with dendrites and dendritic nonlinearities (Hawkins and Ahmad 2016; Schiess, Urbanczik, and Senn 2016; Urbanczik and Senn 2014). These recent models tend to treat the dendritic branch (or sometimes an entire dendritic tree) as discrete loci of nonlinearity. By contrast, the G-clusteron conceptualizes dendritic nonlinearities as being fluid, varying depending on precise synaptic locations. Conceivably, both discrete, branch-dependent nonlinearities and fluid, location-dependent non-linearities could exist simultaneously, further expanding the computational capabilities of real neurons.

From a computational standpoint, we note that the covariance perceptron (Gilson et al. 2019) also takes a correlation-based approach to solving learning tasks.

### Biological Plausibility

The G-clusteron takes several liberties with respect to the biological phenomena from which it is inspired. While a multiplicative activation function may be a tolerable first-order approximation of the synergetic cooperation between NMDA synapses, a sigmoidal function provides a closer fit (Poirazi, Brannon, and Mel 2003). Even to the extent that a multiplicative model does portray the synergistic relationship between excitatory NMDA synapses, such a relationship does not exist between excitatory synapses and inhibitory synapses or between inhibitory synapses and other inhibitory synapses. The multiplicative interaction with negative inputs, however, is essential to the learning protocol of the G-clusteron, so this aspect of our algorithm should be viewed as “biologically inspired” rather than an accurate depiction of what occurs in real biological cells. We do note, however, that clustering dynamics have been observed to occur between dendritic spines and inhibitory synapses (Chen et al. 2012).

The mechanism of learning synaptic locations in the G-clusteron, namely the attraction and repulsion of synapses based on the activity of their presynaptic inputs, also warrants discussion from a biological standpoint. As we have described, the computation of the “forces” exerted by each synapse results in a vector field along the dendrite with regions of attraction and repulsion. Such a vector field could be implemented by the release of attractive or repulsive chemical factors in proportion to the local dendritic activation which diffuse in the extracellular space, creating a distance-dependent gradient that effectively sums the “pull” of the synapses that were active along the dendrite. The attractive factors could stimulate the growth of presynaptic axonal boutons or postsynaptic filopodia at specific regions of the dendrite, while the repulsive factors could eliminate existing synapses.

Brain-derived neurotrophic factor (BDNF) and its precursor, proBDNF, are strong candidates for the signaling mechanism underlying the sort of structural plasticity we suggest here. BDNF has been shown to be responsible for structural plasticity in development by stabilizing correlated synapses during development, while proBDNF weakens synapses that exhibit uncorrelated activity (Niculescu et al. 2018). Another possible signaling agent is estradiol, which is also involved in the formation of new spines (Mendez, Garcia-Segura, and Muller 2011) and interacts with BDNF in spine regulation pathways (Murphy, Cole, and Segal 1998). Astrocytes and microglia, which play an important role in spine elimination (Stein and Zito 2019) may also be implicated in our model as a mechanism for intelligently rearranging synapses on the dendrite in response to local activity.

Several testable experimental predictions emerge from the G-clusteron’s location learning rule. If presynaptic axons are indeed being coaxed into moving their synapses according to chemical gradients, we might expect to see axonal boutons being destroyed and new boutons from the same axon forming a short distance away. We may also expect to see different clustering patterns on different regions of the dendrite due to different lengths constants, as in Figure 2A-B. Because the length constant λ increases with dendritic diameter (Koch 2004; Wilfrid Rall 1959), we might expect that thinner regions of dendrite would have a larger number of small clusters than wider regions, which may tend to group synapses into a smaller number of large clusters. However, these effects would likely depend on the exact nature of the task that the neuron performs.

The weight update rule (Eq. 11) of the G-clusteron can also be understood in a biological framework. Experimental evidence has shown that when LTP is induced in one synaptic spine, the threshold for LTP induction in nearby spines within ∼10µm are reduced (Harvey and Svoboda 2007), a process mediated by the intracellular diffusion of Ras (Harvey et al. 2008). The G-clusteron’s weight update rule qualitatively expresses a similar phenomenon – if a synapse has a strong input such that its weight increases a large amount, nearby synapses will require less input to achieve a large weight update.

### Future Directions

The G-clusteron can be extended in a variety of directions. Because the G-clusteron is inspired by biological dendrites, we use one-dimensional synaptic locations, but our model and algorithm can be easily modified to operate with a “high-dimensional dendrite” where synapses interact with each other as a function of distance in a space with arbitrarily high dimensions, instead of merely moving along a line. Presynaptic axons ostensibly do not just need to localize their boutons within a branch; they may also want to decide between branches of the same neuron or among different neurons. As such, a 2-D or 3-D G-clusteron, where synapses are localized within the plane of a neuron’s branching structure or within a 3D volume of brain tissue, also has some biological motivation.

Our model can also be made more biologically plausible in a number of ways, such as by using a sigmoidal nonlinearity instead of a multiplicative nonlinearity (Poirazi, Brannon, and Mel 2003; Poirazi and Mel 2001), incorporating the most recent work regarding distance-dependent interactions between nonlinear synapses (Behabadi et al. 2012; M. Jadi et al. 2012; M. P. Jadi et al. 2014), or incorporating attenuation from the dendrite to the soma (W Rall 1967; Wilfrid Rall 1959). It would be valuable to see how incorporating these features into the G-clusteron model would affect both the weight and location gradient update rules.

As with any single neuron model, the G-clusteron should also be explored in the context of a multi-layer network. The G-clusteron’s gradient descent-based update rules lend themselves to the possibility of a backpropagation algorithm for a deep network of G-clusteron neurons. This can create exciting avenues for a version of deep learning that incorporates synaptic weight updates together with dendritic nonlinearities and structural plasticity.

## Methods

### Derivation of Location Update Rule for the G-clusteron

We wish to derive a learning rule that updates the synaptic locations on each iteration of the algorithm of the form:

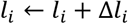

Where Δ*l*_*i*_ is proportional to the gradient of the error with respect to the locations. For an arbitrary loss function J and nonlinearity g where we have J (*g*(*h*(*l*_***i***_))) we have:

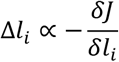

By the chain rule we have:

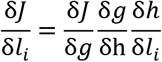

The first two factors of the gradient, 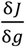 and 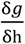, are specific to the cost function and non-linearity chosen. In our case, we use the logit cross-entropy error

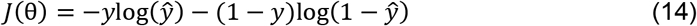

and sigmoidal nonlinearity

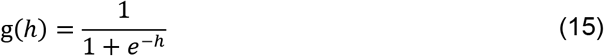

We have:

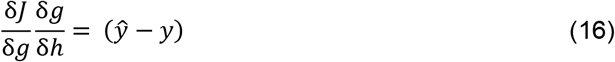

For the final term (see S1 Text for full derivation of 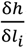):

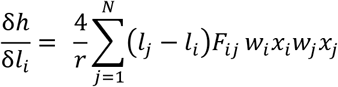

We thus have:

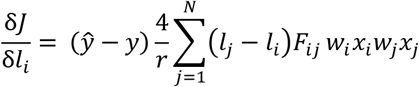

We can subsume the constant factor 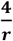 into the learning rate, which gives:

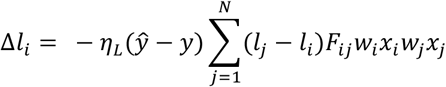

Which gives our location update rule (Eq. 7). This same expression holds if we use a softmax nonlinearity:

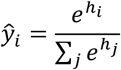

and a cross-entropy cost function:

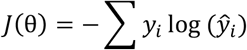

as we do for the multiclass MNIST classification task.

### Derivation of Bias Update Rule

We also require a rule for the update of the bias term *b*. As above:

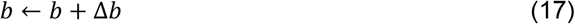

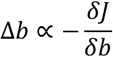

And

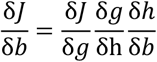

The first two terms are as above, the final term is:

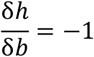

so we have

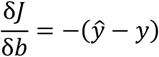

And

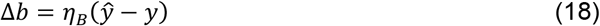

(Note the difference in sign relative to the location and weight rules).

### Derivation of Weight Update Rule

In addition to the location update rule, we also derive a gradient descent rule for the weights of the G-clusteron of the form:

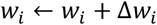

Where η is the learning rate, and:

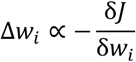

We have

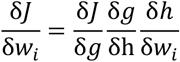

As with the location update rule, we have:

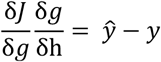

Taking the derivative 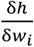 we obtain (see S2 Text for full derivation of 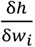):

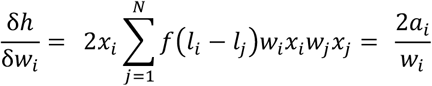

We can subsume the factor 2 into the learning rate. The weight update rule for the G-clusteron is thus:

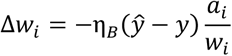

This rule can easily be adapted to a batch learning protocol by simultaneously calculating the activations over the entire batch as shown above.

### Efficient Computation and Batch Learning

For computational efficiency on the MNIST task, the output of the G-clusteron and the learning rules can be implemented in a vectorized fashion. We assume here that we will be working with a dataset comprised of multiple input patterns, running from index *p* ∈ 1,2 … *P*. We will denote input *j* in pattern *p* as 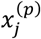. To efficiently calculate each synaptic activation over an entire dataset, we define a matrix *S* such that

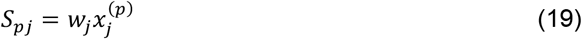

We also define a signed distance matrix *D* where

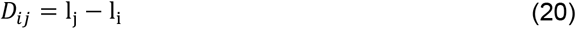

The distance-dependent factor matrix *F* can be expressed in terms of *D*:

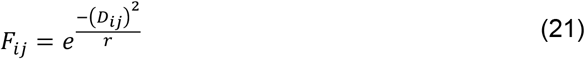

The activation for synapse *j* in pattern *p*, 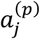, is the element *A*_*pj*_ of matrix *A*, which can be computed as:

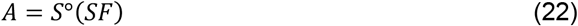

Where ° denotes the Hadamard product, i.e. elementwise multiplication.

For the location learning rule, we utilize a batch protocol where updates are performed after observing *P* patterns (*P* here is the size of the batch, not the dataset). Here, for each pattern *p* in the batch we define a matrix *Q*^(*p*)^ such that

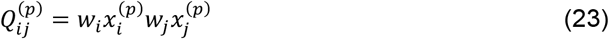

The derivative of the loss function on the entire batch with respect to location *l*_*i*_ is given by:

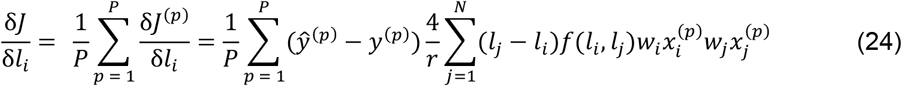

Using our matrix notation, we have

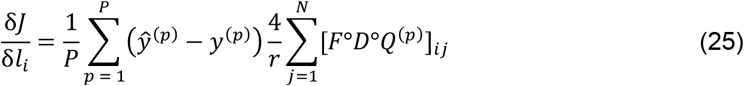

However, this is computationally intensive as it requires an elementwise multiplication for every input pattern in the batch (*F*°*D* can be precomputed for the entire batch, but *F*°*D*°*Q*^(*p*)^ is different for every input pattern). We therefore rearrange and obtain:

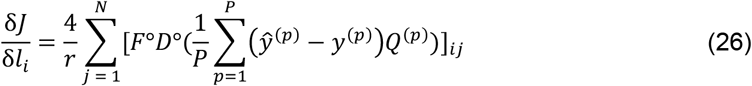

Which averages over all input patterns in the batch first, requiring only two elementwise multiplications regardless of batch size.

### Toy examples

To train the G-clusteron to continually increase or decrease its activation on a particular example or dataset, we treated the activation function as a threshold function which always returned the wrong answer such that it would perform the maximal update at each epoch. For patterns with label *y* = 0, we fixed *ŷ*= 1 such that *ŷ*− *y* = 1, for patterns with label *y* = 1, we fixed *ŷ*= 0 such that *ŷ*− *y* = −1.

**Learning MNIST** (make basically about dataset sizes)

For the MNIST learning tasks, we used the Tensorflow (Abadi et al. 2016) MNIST dataset, which is composed of 60,000 training examples and 10,000 test examples. The examples are split roughly evenly between the 10 digits.

For the one-vs-all experiment, the original clusterons was trained only on positive training examples, ∼6,000 examples per digit. The G-clusterons and logistic regression classifiers were trained on a balanced dataset with ∼6000 images of the digit from the positive class and ∼6000 total images of other digits. The test set was evenly split between positive and negative examples. Hyperparameters for the one-vs-all task can be found in Table 2.

**Table 2.**
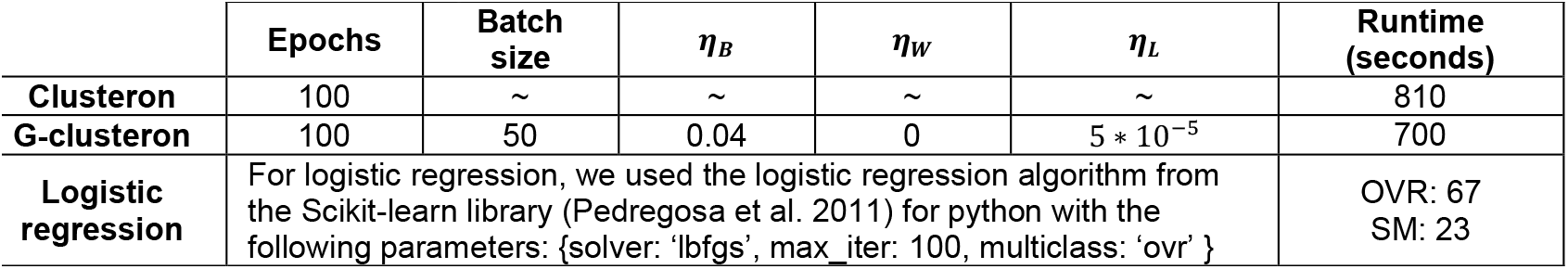
Hyperparameters for one-vs-all learning protocols.

For the all-vs-all experiment, in both the OVR and SM protocols, each of the 10 clusteron nodes was trained on all images from its positive class. Each G-clusteron as well as logistic regression was trained with the entire MNIST dataset. Hyperparameters for all-vs-all are shown in Table 3.

**Table 3.** Hyperparameters for all-vs-all learning protocols. Abbreviations: WR/LR/BR: weight/location/both rules, respectively. OVR: one-vs-rest, SM: softmax.

| | Epochs | Batch size | $\eta_B$ | $\eta_W$ | $\eta_L$ | Runtime (seconds) |
| --- | --- | --- | --- | --- | --- | --- |
| Clusteron | 100 | ~ | ~ | ~ | ~ | 827 |
| G-clusteron (WR, OVR) | 100 | 100 | 0.04 | $10^{-4}$ | 0 | 395 |
| G-clusteron (LR, OVR) | 100 | 100 | 0.04 | 0 | $4 * 10^{-5}$ | 1986 |
| G-clusteron (BR, OVR) | 100 | 100 | 0.04 | $10^{-4}$ | $4 * 10^{-5}$ | 1095 |
| G-clusteron (WR, SM) | 2000 | 30 | $10^{-5}$ | $10^{-5}$ | 0 | 317 |
| G-clusteron (LR, SM) | 2000 | 3 | $5 * 10^{-6}$ | 0 | $5 * 10^{-6}$ | 755 |
| G-clusteron (BR, SM) | 2000 | 5 | $10^{-5}$ | $10^{-5}$ | $10^{-5}$ | 961 |
| Logistic regression | For logistic regression, we used the logistic regression algorithm from the Scikit-learn library (Pedregosa et al. 2011) for python with the following parameters: {solver: 'lbfgs', max_iter: 100, multiclass: 'ovr' for or 'multinomial' for SM} |  |  |  |  | OVR: 67<br>SM: 23 |

For both one-vs-all and all-vs-all (SM and OVR), the hyperparameter *r* of the G-clusteron was always set to 0.23. For the original clusteron, synapses interacted if they were within 10 synapses of each other. Hyperparameters were hand-tuned to maximize accuracy in all cases.

### XOR problem

The XOR function is defined as (Table 4):

**Table 4.** XOR function.

| Input |  | Output |
| --- | --- | --- |
| $x_1$ | $x_2$ | $x_1 \text{ XOR } x_2$ |
| 0 | 0 | 0 |
| 1 | 0 | 1 |
| 0 | 1 | 1 |
| 1 | 1 | 0 |

For the G-clusteron to solve the XOR problem, we require:

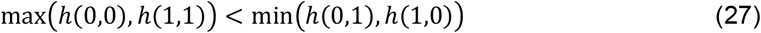

where h(***x***) is the output of the G-clusteron on the input vector ***x*** (In our case, the vector [*x*_1_, *x*_2_]) before applying the sigmoidal nonlinearity, as defined in Eq 4.

In the case of two inputs, h(***x***) can be written as:

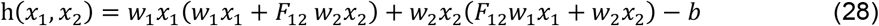

We therefore have (Table 5):

**Table 5.** G-clusteron outputs for two binary inputs.

| Input |  | Output |
| --- | --- | --- |
| $x_1$ | $x_2$ | $h(x_1, x_2)$ |
| 0 | 0 | $-b$ |
| 1 | 0 | $w_1^2 - b$ |
| 0 | 1 | $w_2^2 - b$ |
| 1 | 1 | $w_1^2 + w_2^2 + 2 \cdot F_{12} \cdot w_1 w_2 - b$ |

From Equations 27 and 28 and Table 5, we require:

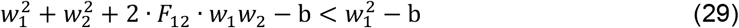

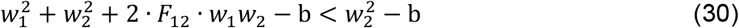

Which can be simplified to Equations 12 and 13 in Results. (Note that there are two other inequalities that follow from Equations 27 and 28, namely that 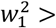 and 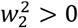, which require that neither *w*_1_ nor *w*_2_ are equal to 0, however Equations 29 and 30 already guarantee this.)

To test the different G-clusteron learning rules on the XOR dataset, we created a G-clusteron for each of the three learning paradigms: weight update rule only, location update rule only, and both weight and location update rules together. Each G-clusteron was run on 1000 trials (i.e. parameter initializations) for 10,000 epochs per trial.

For each trial, the weight values were chosen randomly from a uniform distribution between [-1,1], and the initial locations were chosen in a way such that *F*_12_ was uniformly distributed between [0,1]. The G-clusterons were trained with a stochastic gradient descent protocol such that each epoch, one out of the four input vectors for the XOR function (Table 4) were presented to the algorithm, which would update its parameters according to the update rule. For computational efficiency, we stopped the learning protocol on a given trial when the algorithm converged. Convergence was defined as achieving perfect accuracy on all 4 input vectors of the XOR problem for 10 consecutive epochs. Hyperparameters and the net runtime for all 10,000 trials are shown in Table 6.

**Table 6.**
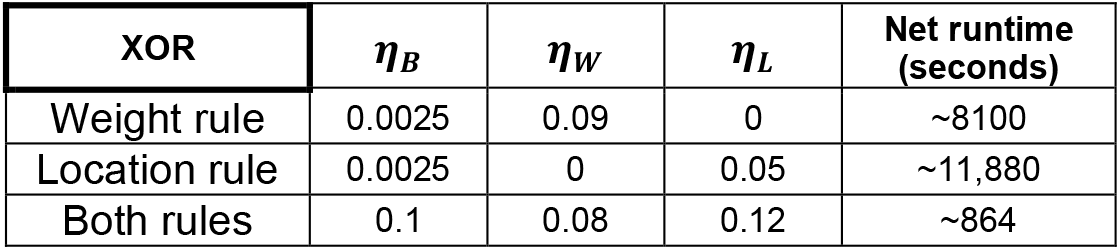
Hyperparameters for XOR problem.

For the XOR problem, the *r* hyperparameter was set to 1. Momentum was not used for the XOR problem. Hyperparameters were hand-tuned to maximize the number of converged trials. Runtimes differed due to different convergence probabilities for the different rules.

All scripts were written in Python and run on a Acer Aspire 5 Notebook laptop with an i7-10510U quad-core processor (1.80 GHz) and 16GB RAM, running on a Windows 10 operating system.

## Supporting Information

**S1 Movie**. (Movie for Figure 2A) Dynamics of synaptic clustering due to the G-clusteron location update rule, using a large value for the *r* parameter and trained a single positive-labeled pattern. Synapses locations are initiated randomly, and then separate into groups of synapses with positive or negative inputs.

**S2 Movie**. (Movie for Figure 2B) Same as S1 Movie but using a small radius instead.

**S3 Movie. (**Movie for Figure 2C) Similar to S1 Movie, but trained on a negative example, with synapse locations initiated such that synapses are clustered with similarly signed input values. With learning, clusters are broken leading to lower overall activation.

**S4 Movie. (**Movie for Figure 3C) Dynamics of synaptic clustering due to the G-clusteron location update rule, using positive labeled patterns from a multivariate gaussian dataset with multiple groups of positively correlated inputs.

**S5 Movie. (**Movie for Figure 3D) Same as S4 Movie, but with negative labeled patterns. Clusters of negatively correlated inputs are formed over the course of learning.

**S1 Text**. Proof that the G-clusteron can approximate a linear classifier by adding an additional “bias synapse” and setting the weights and bias appropriately.

**S2 Text**. Extended derivation of location update rule.

**S3 Text**. Extended derivation of weight update rule.

## Acknowledgements

The authors would like to thank Eyal Gal and Itamar Landau for their contributions to the development of the G-Clusteron plasticity rules. We would also like to thank David Beniaguev for some helpful suggestions and Ilenna Jones for her valuable comments on an early version of this manuscript.

