## Supplementary material for "The Gradient Clusteron: A model neuron that learns via dendritic nonlinearities, structural plasticity, and gradient descent": S1 Text

**S1. Proof that the G-clusteron can approximate a linear classifier**

To prove that every linear classifier of the form $h(x) = \sum_{i=1}^{N} w_i x_i - b$ can be approximated to arbitrary precision by a weighted G-clusteron (with any arrangement of synaptic locations) of the form $h^*(x) = \sum_{i=0}^{N} w_i^* x_i \sum_{j=0}^{N} F_{ij} w_j^* x_j - b^*$ with the inclusion of a single "bias synapse" $x_0$ ($x_0 = 1$ for all input patterns, this is distinct from the zeroth-order bias term $b^*$), we observe that:

$$h^*(x) = w_0^{*2} x_0^2 + \sum_{i=1}^{N} 2F_{i0} w_i^* x_i w_0^* x_0 + \sum_{i=1}^{N} w_i^* x_i \sum_{j=1}^{N} F_{ij} w_j^* x_j - b^* \qquad \text{(S1.1)}$$

Because $x_0 = 1$, this expression simplifies to:

$$h^*(x) = w_0^{*2} + \sum_{i=1}^{N} 2F_{i0} w_i^* x_i w_0^* + \sum_{i=1}^{N} w_i^* x_i \sum_{j=1}^{N} F_{ij} w_j^* x_j - b^* \qquad \text{(S1.2)}$$

We set the weights of the G-clusteron (other than the bias synapse weight $w_0^*$):

$$w_i^* = \frac{w_i}{2F_{i0} w_0^*} \; for \; i > 1 \qquad \text{(S1.3)}$$

And we set the bias of the G-clusteron as:

$$b^* = w_0^{*2} + b \qquad \text{(S1.4)}$$

We therefore have:

$$\begin{aligned} h^*(x) &= \sum_{i=1}^{N} w_i x_i + \sum_{i=1}^{N} \frac{w_i}{2F_{i0} w_0^*} x_i \sum_{j=1}^{N} \frac{F_{ij} w_j}{2F_{j0} w_0^*} x_j - b \\ &= \sum_{i=1}^{N} w_i x_i - b + \frac{1}{w_0^{*2}} \sum_{i=1,j=1}^{N} \frac{F_{ij} w_i w_j}{4F_{i0} F_{j0}} x_i x_j \end{aligned} \qquad \text{(S1.5)}$$

If we set $w_0^*$ to be a large number (greater than 1) such that:

$$w_0^* \gg \max_{i,j \in [1 \ldots N]} \left( \frac{|w_i w_j|}{4F_{i0} F_{j0}} \right) \qquad \text{(S1.6)}$$

The final term for equation (S1.5) approaches $0$, and we therefore have:

$$h(x) \approx \sum_{i=1}^{N} w_i x_i - b \qquad \text{(S1.7)}$$

Note that this approximation can be obtained for any arrangement of the synaptic locations in the G-clusteron as long as the weights are appropriately set as in equation S1.3.
