## Supplementary material for "The Gradient Clusteron: A model neuron that learns via dendritic nonlinearities, structural plasticity, and gradient descent": S2 Text

**S2. Extended Derivation of Location Update Rule**

To derive the location update rule, we need to find the derivative of the raw output of the G-clusteron $h$ with respect to an arbitrary location $l_k$. (In Results and Methods we used the notation $\frac{\delta h}{\delta l_i}$; here we use the notation $\frac{\delta h}{\delta l_k}$ to avoid confusion with the summation indices):

$$\frac{\delta h}{\delta l_k} = \frac{\delta}{\delta l_k}\left(\sum_{i=1}^{N} a_i - b\right) \tag{S2.1}$$

$$= \frac{\delta}{\delta l_k}\left(\sum_{i=1}^{N} w_i x_i \sum_{j=1}^{N} e^{\frac{-\left(l_i - l_j\right)^2}{r}} w_j x_j - b\right) \tag{S2.2}$$

We break this expression into four terms reflecting the four possible conditions: $(i = k, j = k), (i \neq k, j \neq k), (i = k, j \neq k), (i \neq k, j = k)$

$$= \frac{\delta}{\delta l_k}\left(w_k x_k w_k x_k + \sum_{i=1, i\neq k}^{N} w_i x_i \sum_{j=1, j\neq k}^{N} e^{\frac{-\left(l_i - l_j\right)^2}{r}} w_j x_j + w_k x_k \sum_{j=1, j\neq k}^{N} e^{\frac{-\left(l_k - l_j\right)^2}{r}} w_j x_j + \sum_{i=1, i\neq k}^{N} w_i x_i\, e^{\frac{-(l_i - l_k)^2}{r}} w_k x_k - b\right) \tag{S2.3}$$

We calculate the derivative of each term separately:

| Term | $\frac{\delta}{\delta l_k}$ |
|---|---|
| $w_k x_k w_k x_k$ | 0 |
| $\sum_{i=1, i\neq k}^{N} w_i x_i \sum_{j=1, j\neq k}^{N} e^{\frac{-\left(l_i - l_j\right)^2}{r}} w_j x_j$ | 0 |
| $w_k x_k \sum_{j=1, j\neq k}^{N} e^{\frac{-\left(l_k - l_j\right)^2}{r}} w_j x_j$ | $-\frac{2}{r} w_k x_k \sum_{j=1, j\neq k}^{N} (l_k - l_j) e^{\frac{-\left(l_k - l_j\right)^2}{r}} w_j x_j$ |
| $\sum_{i=1, i\neq k}^{N} w_i x_i\, e^{\frac{-(l_i - l_k)^2}{r}} w_k x_k$ | $\frac{2}{r} \sum_{i=1, i\neq k}^{N} w_i x_i\, (l_i - l_k) e^{\frac{-(l_i - l_k)^2}{r}} w_k x_k$ |

Noting that the expressions in the last two rows are equivalent, summing the derivatives of all four terms gives us:

$$\frac{4}{r} w_k x_k \sum_{j=1, j\neq k}^{N} w_j x_j \left(l_j - l_k\right) e^{\frac{-\left(l_j - l_k\right)^2}{r}} \tag{S2.4}$$

Noting that when $j = k$, $w_j x_j \left(l_j - l_k\right) e^{\frac{-\left(l_j - l_k\right)^2}{r}} = 0$, we have:

$$\frac{\delta h}{\delta l_k} = \frac{4}{r} \sum_{j=1}^{N} \left(l_j - l_k\right) e^{\frac{-\left(l_j - l_k\right)^2}{r}} w_k x_k w_j x_j \tag{S2.5}$$
