## Supplementary material for "The Gradient Clusteron: A model neuron that learns via dendritic nonlinearities, structural plasticity, and gradient descent": S3 Text

**S3 - Extended Derivation of Weight Update Rule**

As with the location rule (eq. S1.1), we need to find the derivative of the raw output of the G-clusteron $h$ with respect to an arbitrary weight $w_k$:

$$\frac{\delta h}{\delta w_k} = \frac{\delta}{\delta w_k}\left(\sum_{i=1}^{N} a_i - b\right) \tag{S3.1}$$

As with the location rule, we break $h$ up into four terms and calculate the derivatives separately:

| Term | $\frac{\delta}{\delta w_k}$ |
|---|---|
| $w_k x_k w_k x_k$ | $2x_k^2 w_k$ |
| $\sum_{i=1,i\neq k}^{N} w_i x_i \sum_{j=1,j\neq k}^{N} e^{\frac{-(l_i-l_j)^2}{r}} w_j x_j$ | 0 |
| $w_k x_k \sum_{j=1,j\neq k}^{N} e^{\frac{-(l_k-l_j)^2}{r}} w_j x_j$ | $x_k \sum_{j=1,j\neq k}^{N} e^{\frac{-(l_k-l_j)^2}{r}} w_j x_j$ |
| $\sum_{i=1,i\neq k}^{N} w_i x_i\, e^{\frac{-(l_i-l_k)^2}{r}} w_k x_k$ | $\sum_{i=1,i\neq k}^{N} w_i x_i\, e^{\frac{-(l_i-l_k)^2}{r}} x_k$ |

Noting that the expressions in the last two rows are equivalent, summing the derivatives of all four terms gives us:

$$= 2x_k^2 w_k + \sum_{j=1,j\neq k}^{N} 2e^{\frac{-(l_k-l_j)^2}{r}} w_j x_j x_k \tag{S3.2}$$

Noting that when $j = k$:

$$2e^{\frac{-(l_k-l_j)^2}{r}} w_j x_j x_k = 2x_k^2 w_k \tag{S3.3}$$

We have:

$$\frac{\delta h}{\delta w_k} = 2x_k \sum_{j=1}^{N} e^{\frac{-(l_k-l_j)^2}{r}} w_j x_j = 2\frac{a_k}{w_k} \tag{S3.4}$$
